# Exploring the conformational ensembles of protein-protein complex with transformer-based generative model

**DOI:** 10.1101/2024.02.24.581708

**Authors:** Jianmin Wang, Xun Wang, Yanyi Chu, Chunyan Li, Xue Li, Xiangyu Meng, Yitian Fang, Kyoung Tai No, Jiashun Mao, Xiangxiang Zeng

## Abstract

Protein-protein interactions are the basis of many protein functions, and understanding the contact and conformational changes of protein-protein interactions is crucial for linking protein structure to biological function. Although difficult to detect experimentally, molecular dynamics (MD) simulations are widely used to study the conformational ensembles and dynamics of protein-protein complexes, but there are significant limitations in sampling efficiency and computational costs. In this study, a generative neural network was trained on protein-protein complex conformations obtained from molecular simulations to directly generate novel conformations with physical realism. We demonstrated the use of a deep learning model based on the transformer architecture to explore the conformational ensembles of protein-protein complexes through MD simulations. The results showed that the learned latent space can be used to generate unsampled conformations of protein-protein complexes for obtaining new conformations complementing pre-existing ones, which can be used as an exploratory tool for the analysis and enhancement of molecular simulations of protein-protein complexes.

## INTRODUCTION

Proteins are the basic components of life, and almost all proteins perform their functions through non-covalent interactions with proteins, peptides, nucleotides, and small molecules, and so on^1,2^. Protein-protein interactions (PPIs) play a key role in many biological processes. Understanding the 3D structure of protein-protein complexes can help to understand many biological processes and molecular mechanisms, providing useful insights for drug and protein design^2–7^. The number of protein-protein complex structures that were determined by experimental techniques such as X-ray crystallography and cryo-electron microscopy (cryo-EM) is increasing. However, obtaining these experimental structures is still quite time-consuming and costly, and high-resolution structures determination and dynamic mechanism measurements are still experimentally challenging. Computational modeling of protein-protein complexes based on limited experimental data is a helpful complement and alternative to limited experimental approaches. Traditional approaches consist of protein-protein docking and homology modeling^8–10^. Deep learning (DL) has recently made unprecedented advances in the field of computational biology^11^, in particular the various applications of AlphaFold2 ^12^,AlphaFold-Multimer^6^ and RoseTTAFold^13^ for the prediction of protein structures and protein-protein complexes. Unfortunately, experimental structures usually only capture static conformations in the low-energy state for protein-protein binding. Although molecular modeling or DL have been applied to predict protein-protein complex structures, their sampling of stable functional conformations, protein binding kinetics, and conformational space remains challenging^2–4,14–16^.

Molecular dynamics (MD) simulations have wide applications for the conformations and dynamics of proteins and protein-protein complexes^2,3,14,17–19^. MD samples from potential conformations of biomolecular systems to determine the most energetically favorable subspace in the conformational space. These simulations have greatly benefited from hardware and software developments such as graphics processing unit (GPU), AMBER^20,21^, GROMACS^22^, and OpenMM^23^. Although these advances have been made, MD simulations to produce dynamic conformational ensembles remain challenging in terms of computational cost and efficient sampling of the conformational ensembles. Protein-protein binding affinity is often stronger, and the process of binding and unbinding often requires longer time scales^3^. Although many MD simulations and enhanced sampling methods have been developed for PPIs modeling, the conformational ensembles of protein-protein complex remain largely underexplored.

Deep learning and generative artificial intelligence have recently made unprecedented advances in computational biology^11,24,25^. Generative deep learning has been successful for the sampling of protein conformations^25^, such as long short-term memory (LSTM)^26^, autoencoders (AEs)^27,28^, variational autoencoders (VAEs)^29–33^, generative adversarial networks (GANs)^31,34^, score-based models^35^, energy-based models^36^, Transformer^31,37,38^, and active learning^33^. Training of deep generative models relies on conformations extracted from molecular dynamics simulations. Aims to learn the data distribution of molecular dynamics simulations is used to generate novel realistic ensembles of protein conformations^2,4,14^.

In this study, we proposed a Transformer-based generative framework to accelerate the sampling and exploration of the conformational ensembles of protein-protein complexes. First, the crystal structure of the barnase-barstar complex was performed in 100 ns MD production simulations. The iterations were repeated to obtain six trajectories meeting specific criteria. Then the trajectories data were split into 300 ensembles (1000 frame/ensemble) as a training set. We use the Transformer-based generative neural network to learn the 3D conformational distribution of protein-protein complexes in the training set. Our model is named AlphaPPImd, which directly generates novel conformations of protein-protein complex and generates conformations beyond the MD time scale. The results of scoring the generated conformational models of protein-protein complexes show that most of the generated conformational models were acceptable models. Conformational ensembles analysis shows that the learned latent space can be used to generate unsampled conformations of protein-protein complexes. We visualize the key amino acid residues and residue pairs of the protein-protein complexes with higher weights captured by the attention mechanism of the model. The results show that the deep generative model learns key residues from multiple MD trajectories that influence the conformational and dynamical mechanisms of protein-protein complexes, provides mechanistic insights into protein-protein binding. Generative deep learning can be utilized as one of the optional tools to enhance the exploration of the conformational ensembles of protein-protein complexes and the analysis of key amino acid residues.

## MATERIALS AND METHODS

### Molecular Dynamics Simulations

The crystal structure of the barnase-barstar complex (PDB ID: 1BRS)^39^ was used to build simulation system and has been widely used in previous research^3,14^. We downloaded the crystal structure from the Protein Data Bank (PDB)^40,41^, and extracted chain A and chain D as the initial complex structure by removing the ligand and crystal water. The missing hydrogen atoms were added by tleap module in AmberTools^42^. The system was neutralized by adding Na^+^ and Cl^-^ ions, and solvated in a 12 angstrom (Å) periodic bounding box of TIP3P water molecules^43^. The topology and coordinate files of the system were prepared using the tleap module in AmberTools and the AMBER ff14SB force field^44^.

The simulated system was first energy minimized by performing a 500 steps typical ensemble (NVT) simulation with the Langevin integrator, and then further equilibrated by performing a 10,000 steps NPT simulation at 300 K. Particle Mesh Ewald (PME) algorithm was applied to the calculation of long-range electrostatic interactions^45^. A cutoff 1 nm for direct space interactions was used and the SHAKE algorithm was set to constrain the lengths of all bonds involving hydrogen atoms^46^. The simulation time step was set to 2 fs, and then six independent 100 ns MD production were performed. All simulations were conducted by OpenMM 7.7^23^.

### Model Architecture of AlphaPPImd

In this section, we introduce the Transformer-based AlphaPPImd framework. The approach is inspired by a recent research in exploring the protein conformational ensembles^31^, which utilized deep generative models to capture the conformational states of protein that are difficult to analyze by traditional molecular dynamics. We first prepared MD trajectories of protein-protein complex. Then, to embed each frame of the protein-protein complex, we constructed two encoders to embed the trajectory features of each chain of the protein-protein complex separately. Explainability is key to attention-based models, and we calculated the weights between residue pairs in the conformational ensembles of protein-protein complex. Finally, to make it possible to recognize the complex and reconstruct the 3D structure of the protein-protein conformation, the protein-protein 3D structures generated by the decoder were aligned for chain and amino acid numbers, and complex structure optimization.

According to the important role of the attention mechanism in the Transformer^47^, Fernández proposed a frame-to-frame Transformer based on the extension of the basin-encoded trajectories^48^. In this study, we are concerned with converting the basin-encoded frame n (the source) of protein-protein complex to the basin-encoded frame n+1 (the target) of protein-protein complex.

The core of the AlphaPPImd framework is self-attention mechanism, which can capture key amino acid residue pairs that affect the conformation of protein-protein complex from MD trajectories. AlphaPPImd consists of the following modules (**Figure 1**). First, the MD trajectories of the protein-protein complex are preprocessed to get sequence length, sequence composition and amino acid residue types for both chains. The torsional *ϕ*, *ψ* angles of selected residues in the trajectories are calculated to represent the conformational state as a string of occupancies in the four basins indicated by the *ϕ*, *ψ* plot^31,48^. Second, the embedding module adds two positional embeddings to the amino acid sequence embedding to generate the frame embedding. Two positional embeddings, correlating the basins of basin-encoded frame with their relative positions in the protein sequence, and correlating the basins of basin-encoded frame with the types of amino acids at each position. Then, each frame embedding of protein-protein complex MD trajectories via the embedding module was input to encoder module of AlphaPPImd, which contains the multi-head self-attention mechanism, attention score, and feature optimization module. To represent the conformational information of the protein-protein complex, we used two encoder modules to capture the conformational state of each of the two chains. Next, the decoder module of AlphaPPImd is designed to learn and capture the cooperative contribution of different types and positions of residues of protein-protein complex to protein-protein conformation. The decoder module mainly consists of masked multi-head self-attention layers, layer normalization layers and feedforward layers, and then apply dropout technique to enhance the robustness. Finally, the prediction module iteratively generates the basins for next frame, and then the Modeller (v10.2) reconstructed protein-protein complex conformational model derived from the extended basin-encoded trajectory^49,50^.

**Figure 1.**
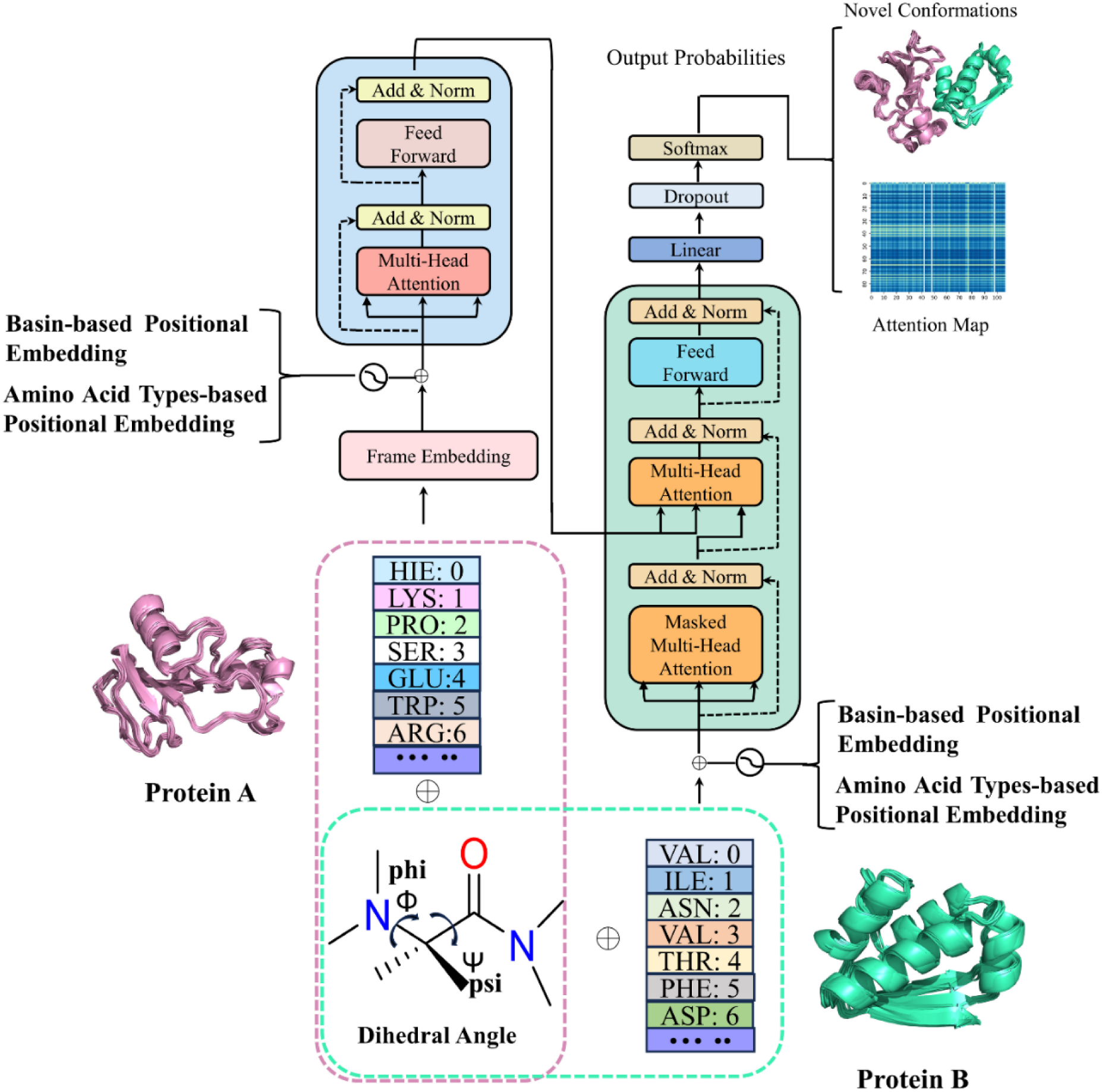
Overview of the AlphaPPImd architecture. The Transformer input is the basin-encoded frames of protein-protein complex. The torsion *ϕ*, *ψ* angles of the all residues of the protein-protein complex are required to be encoded, and then the conformational state of the protein-protein complex is represented as cooperative contribution to the protein-complex conformation indicated by *ϕ*, *ψ*. Multiple short MD trajectories in atomic detail are utilized to train Transformer to propagate protein-protein complex dynamics beyond the time scales accessible to MD. The Transformer outputs novel states of protein complex indicated by *ϕ*, *ψ* that are decoded as conformational model via Modeller^49,50^.

### Attention Mechanism of AlphaPPImd

Attention mechanism is widely used to regulate the weights of feature vectors. The multi-head self-attention layer of the AlphaPPImd decoder module learns the interactions between specific residue pairs. The attention function can be thought of as a mapping between a query (Q) and the output of a key-value pairs (K-V). AlphaPPImd takes the residue embedding of protein-protein complex as Q, regards the global protein-protein complex features as K and V, and calculates the attention weight by using Q and K. The calculation formula is as follows:

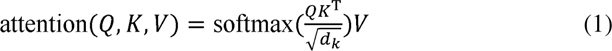

The attention layer updates the values of the statistical features based on the usefulness of critical residue pairs for the conformation of protein-protein complex.

### Model Training and Hyperparameters

The six independent 100 ns MD trajectories of the barnase-barstar complex were split into 300 ensembles, each ensemble consisting of 1000 frames. MD trajectories are preprocessed to keep only protein atoms. Each MD runs contributes a limited set of physical snapshots of protein-protein complex^48^. Each frame in the trajectory was represented as a basins encoded by Φ, Ψ indication. Thus, the torsional state of the protein-protein complex is reduced to a textual representation that retains the dominant subordinate features of the dynamics. All the trajectory ensembles are used as training sets, the validation set is always the trajectory with index 0 and the test set is always the trajectory with index 8.

Model training was conducted on Ubuntu Linux 20.04.6 LTS system. The CPU is an Intel(R) Xeon(R) Gold 6226R CPU@ 2.90 GHz. The memory is 376 GB. The model was trained on the single NVIDIA RTX A5000 GPU, the code language is Python 3.7.12, and the model was constructed by TensorFlow 2.8.0 and Keras 2.8.0. The training consists of 10 epochs and each epoch contains 300 subepochs.

To optimize the model, we use the RMSprop optimizer (learning_rate=0.001), and the loss function is sparse_categorical_crossentropy, and the metrics is accuracy. The number of trainable parameters for each component of the Transformer model in this study can be found in **Supporting Table S1**.

### Evaluation Metrics

The following metrics were used in this study to assess model performance: accuracy, root mean square deviation (RMSD), and a quality measure for protein-protein complex models (DockQ)^51^.

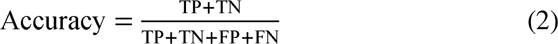

where TP is true positive, TN is true negative, FP is false positive, and FN is false negative.

Root-mean-square deviation (RMSD). RMSD was utilized to quantify conformational differences between the trained and decoded conformations for protein-protein complex. When performing a comparison between the conformation of the generated protein-protein complex and the reference crystal structure, the lower the RMSD, the more accurately the generated conformation; RMSD < 2 Å is generally considered to be very close to the crystal structure conformation, instead, high RMSD levels indicates an inaccurate generated conformation.

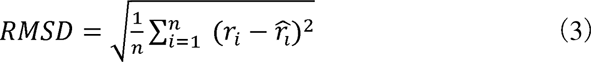

We evaluate the accuracy of the generated conformations of protein-protein complex by DockQ score^51^, a quality score of protein-protein models widely used in the computational structural biology community.

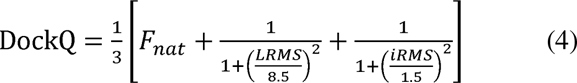

DockQ is a score between 0 and 1, presented based on all the evaluation criteria used in the Critical Assessment of Prediction Interaction (CAPRI). DockQ score is classified as follows: “incorrect”(0.00 ≤ DockQ < 0.23), “acceptable quality” (0.23 ≤ DockQ < 0.49), “medium quality” (0.49 ≤ DockQ < 0.80) and “high quality” (DockQ ≥ 0.80).

Novelty: If the Euclidean distance of the backbone between the generated correct conformations (DockQ ≥ 0.23) of protein-protein complex and the MD conformations in the training set is >2, the generated correct conformations are considered to be novel. The formula of calculating the Euclidean distance is as follows:

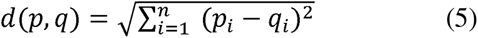

where *p* and *q* represent the generated conformations of protein-protein complex and MD conformations in the training set, respectively, and *p_i_* and *q_i_* are the coordinates of the *i*-th residue of the backbone.

## RESULTS AND DISCUSSION

### MD Trajectories Representation of PPI complex

The RMSD distributions of protein-protein complex for six independent 100 ns MD simulations are shown in **Supporting Figure S1**. The barnase-barstar complex in the training ensembles of the MD trajectories consists of two different chains and 197 residues (barnase chain: 108 residues, barstar chain: 89 residues), and thus only 195 pairs (barnase chain: 107 pairs, barstar chain: 88 pairs) of *ϕ*, *ψ*dihedral angles can be estimated for every time step of the trajectories. As shown in **Figure 2a and 2b**, the Ramachandran plot of the torsional information of the barnase and barstar chain from an ensemble of MD trajectories used in this study, which contains 1000 frames. With each point in the plot representing the backbone torsional state of a residue in one of the frames in the trajectory.

**Figure 2.**
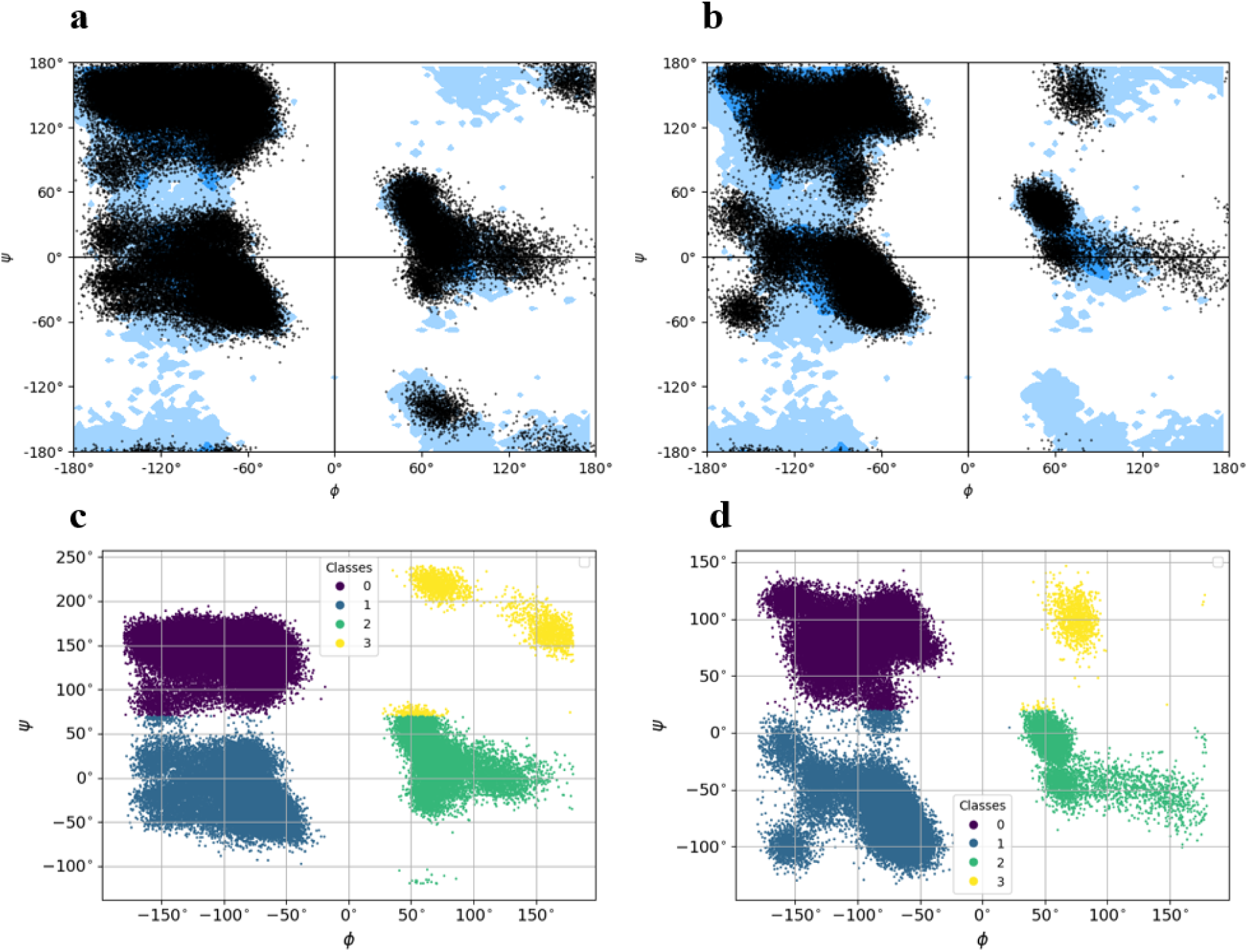
**a**, Ramachandran plot (allowed: dark blue, marginally allowed: lighter blue) of the first trajectory in the barnase chain set. Each point in the plot is a *ϕ*, *ψ* pair of one residue in one frame of the trajectory. **b**, Ramachandran plot (allowed: dark blue, marginally allowed: lighter blue) of the first trajectory in the barstar chain set. Each point in the plot is a *ϕ*, *ψ*pair of one residue in one frame of the trajectory. **c**, Classification of the points in the Ramachandran plot as belonging to 1 of 4 basins in the barnase chain. **d**, Classification of the points in the Ramachandran plot as belonging to 1 of 4 basins in the barstar chain.

The points of the plot were classified into 4 clusters (basins) labeled 0, 1, 2, and 3 by the KMeans algorithm (**Figure 2c and 2d**). The centroids of each cluster are recorded and stored for reconstruction of the full atomistic model of the barnase-barstar complex from the basin-encoded torsional states. Then, each frame of trajectories was converted into a character vector of an alphabet consisting of 4 symbols corresponding to 4 clusters. A similar representation procedure was performed for all 300 ensembles in the MD trajectories dataset for the barnase-barstar complex.

In summary, the barnase-barstar complex is a heterodimer. There are distinctly different residue basins encoded in the two chains. Thus, implying that there are significant differences in generating novel basin-encoded frames and reconstructing conformational models with individual proteins.

### Model Performance and Conformational Reconstruction

The trajectory ensemble datasets of barnase-barstar complex were selected as the datasets. The model was trained with 300 ensembles of the barnase-barstar complex, each ensemble contains 1000 frames. The accuracy and loss of the Transformer model of the barnase-barstar complex obtained by the training and validation sets of the basin-encoded trajectories are shown in **Figure 3a and 3b**. The average training accuracy of the model is 0.995 and the average validation accuracy is 0.999. Although the AlphaPPImd soon achieved stable performance, to further refine the Transformer model and enrich the MD conformational distribution learned by the model, we used multiple MD trajectories as a dataset. Rapid convergence of the sequence that remains the same in subsequent expansion cycles can be avoided by introducing entropy in the frame prediction module.

**Figure 3.**
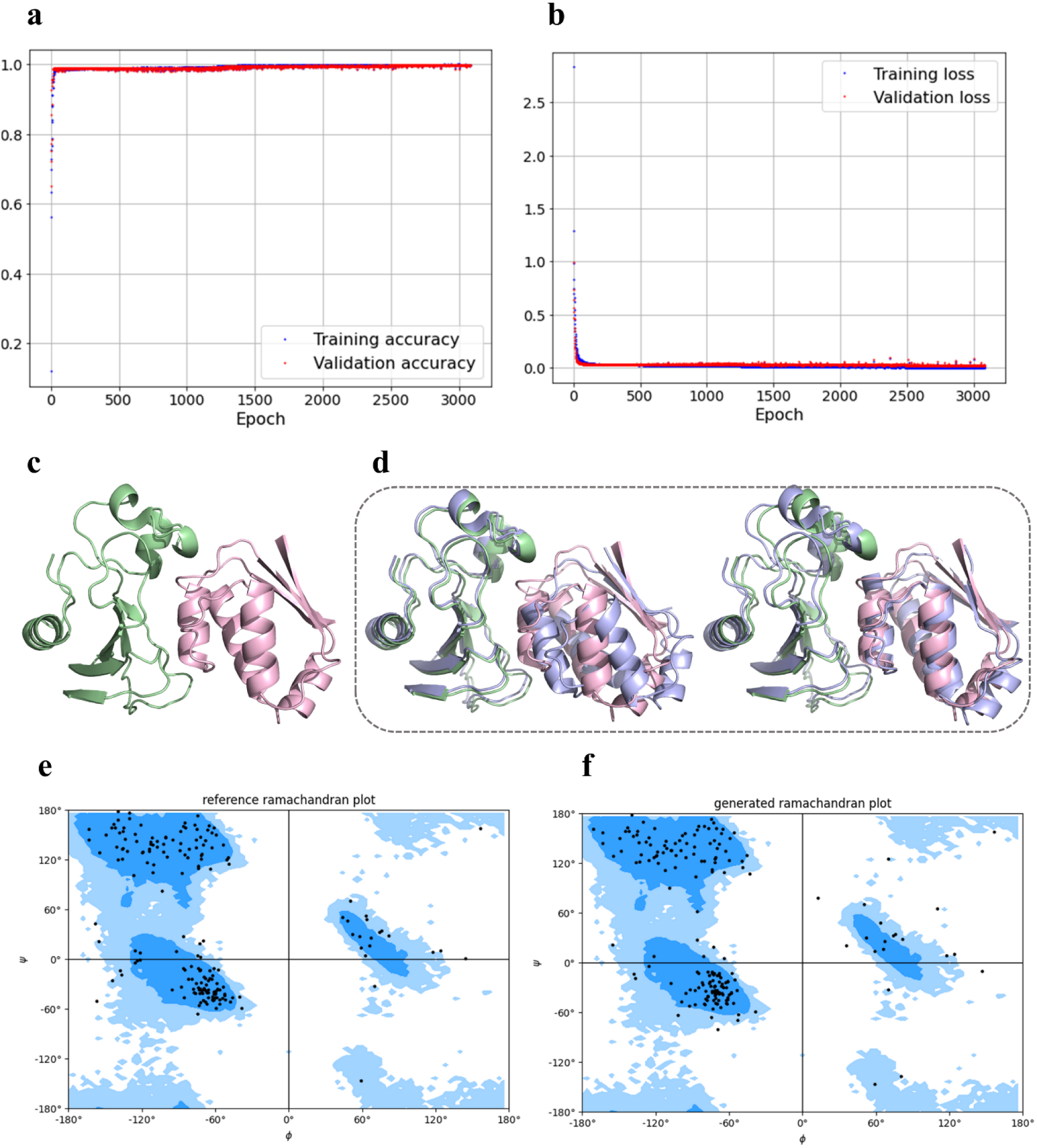
**a and b**, Training and validation set accuracy and loss for the Transformer neural networks. **c**, The conformation of the barnase-barstar complex in one frame of the test sets. **d**, Overlap of the conformation of the generated barnase-barstar complex (color: lightblue) with the reference conformation. **e**, Ramachandran plot of reference conformation. **f**, Ramachandran plot of generated conformation.

A random frame from trajectory of the test set was taken as input, and 100 basin-encoded frames were generated by the trained AlphaPPImd framework. The barnase-barstar complex conformation of the input frame is shown as in **Figure 3c**. The generated extended basin-encoded frames were reconstructed to obtain the conformational model by Modeller^49,50^. The overlap models of the generated conformation and the input conformation are shown in **Figure 3d**. The results show the ability of the model in successfully sampling and expanding the conformations. The Ramachandran plot of the reference structure of the barnase-barstar complex is shown in **Figure 3e**. The Ramachandran plot of the generated conformational model with Φ, Ψ constraints derived by using the center point is shown in **Figure 3f**. The comparison of Figure 3e and 3f shows that the overall conformation of the reference structure is preserved in the corresponding generated conformation. The results show that Modeller correctly performs the dihedral restraints.

### Evaluation of Generated Conformational Ensembles

The MD trajectories of the barnase-barstar complex were randomly selected 1000 frames input into the model to generate 1000 novel basin-encoded frames, which were then reconstructed as novel models of the barnase-barstar complex. The RMSD distribution of 1000 frames of conformations randomly selected from the MD trajectories of the barnase-barstar complex is shown in **Figure 4a**, and the RMSD distribution of 1000 conformations of the barnase-barstar complex that are generated by the AlphaPPImd is illustrated in **Figure 4b**. The RMSD values obtained by comparing the conformations in the MD ensembles of the barnase-barstar complex with the reference crystal structure are all less than 2 Å (**Figure 4a**). The RMSD values obtained by comparing the conformations in the generated ensembles of barnase-barstar complex with the reference crystal structure are mostly less than 2Å (**Figure 4b**). The result reveals that the AlphaPPImd framework generates novel conformational models of protein-protein complex mostly very close to the crystal structure.

**Figure 4.**
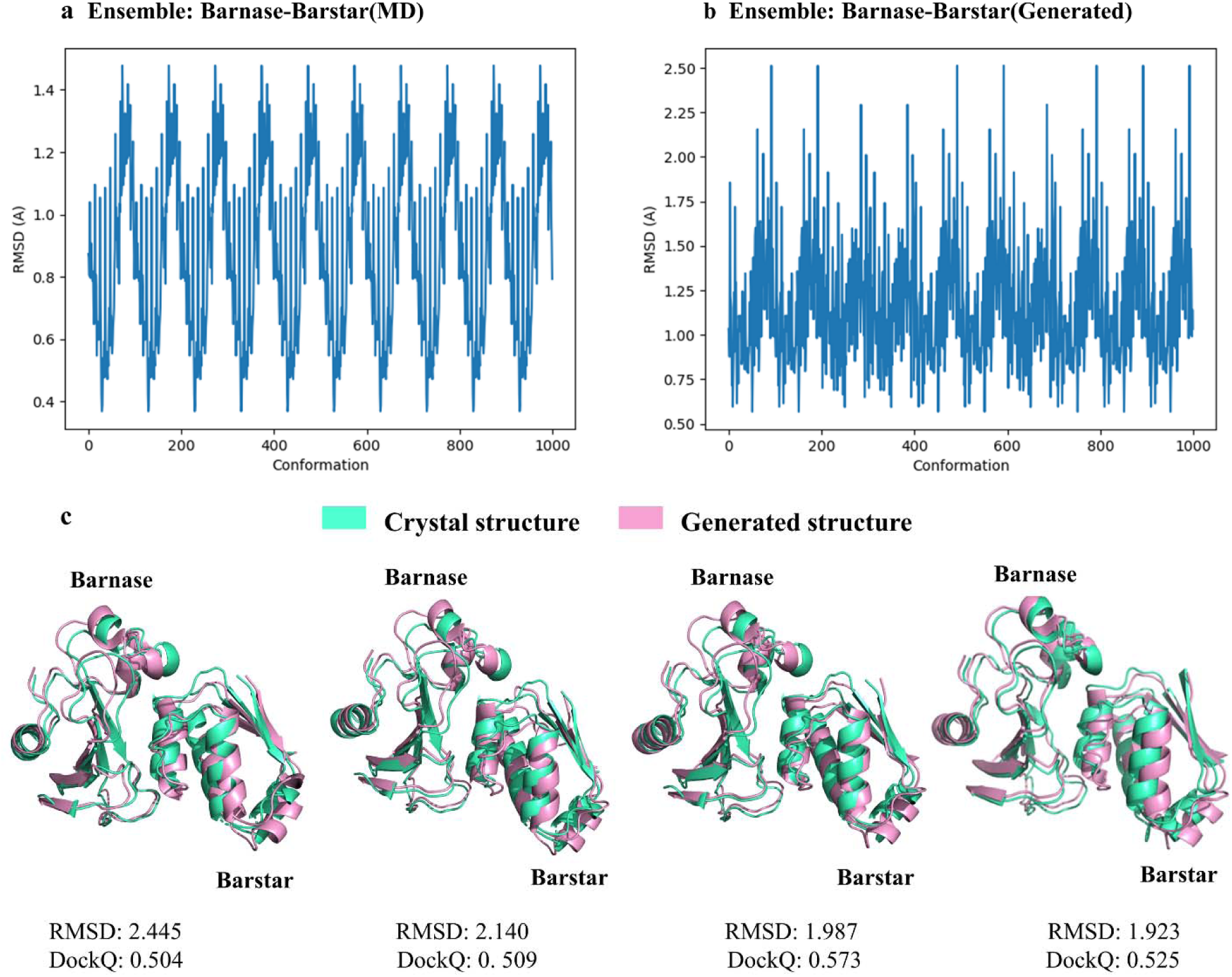
The RMSD distribution of protein-protein complex conformations. **a**, the RMSD distribution of 1000 frames of the barnase-barstar complex MD trajectories. **b**, the RMSD distribution of 1000 conformations of the barnase-barstar complex generated by AlphaPPImd. **c**, overlap of the generated conformational models of the barnase-barstar complex (color: pink) with the crystal structure of the barnase-barstar complex (color: greencyan), show the RMSD values and DockQ score of the generated conformational models with the crystal structure.

We selected four representative conformations with RMSD close to 2 Å from the 1000 conformational models of barnase-barstar complexes generated by the AlphaPPImd model. Then, they were overlapped with the crystal structure, and the RMSD and DockQ score were calculated as shown in **Figure 4c**. As the RMSD becomes smaller, the DockQ score of the generated conformations of the barnase-barstar complex is becoming larger, meaning that the novel protein-protein complex conformational models generated by AlphaPPImd are closer to the reference crystal structure (**Figure 4c**), with higher accuracy (RMSD < 2 Å) and acceptability (DockQ ≥ 0.23). Most of the protein-protein complex conformational models generated by the deep generative model were acceptable (RMSD < 2 Å and DockQ ≥ 0.23). When performing a comparison the conformational ensembles of protein-protein complex MD with the conformational ensembles of generated protein-protein complex, the generated conformations of protein-protein complex had lower RMSD and DockQ scores. A possible reason is that the basin-encoded frame is a simplified textual representation.

### Conformational Space Analysis

The principal component analysis (PCA) is widely used for understanding the dynamics of biological systems^52^. For generating PCA data for ensembles, each frame of the conformational models in the ensembles are overlapped on the initial crystal structure. Then features of each conformational models are calculated for the projection in two-dimensional space. The conformational structures of 1000 randomly selected frames from the MD trajectories of the barnase-barstar complex and 1000 generated structures by the AlphaPPImd model were performed by PCA via ProDy^53^. As shown in **Figure 5a**. The 1000 frame conformational models generated by the AlphaPPImd model were filtered with DockQ < 0.23 (“incorrect”), and then PCA analysis was performed with conformational models extracted from the MD trajectories (**Figure 5b**). The visualization of the Euclidean distance of the backbone between the correct conformations of the generated protein-protein complex (DockQ ≥ 0.23) and the MD conformations in the training set is shown in Supporting Figure S2. The MD systems and the sampling ensembles share the same conformational space. The sampling ensembles of the deep generative model covers the MD trajectory ensembles, and samples novel conformations beyond the MD time scale. Although there are a few incorrect (DockQ < 0.23) conformational models in the conformations generated by the deep generative model, it generates mostly acceptable (DockQ ≥ 0.23) and diverse conformations (**Supporting Figure S2**). The comparison was performed between the generated incorrect conformations and the generated correct conformations, and the visualization of the conformational models shows that the incorrect conformations have more distance between the two chains and lacks interface close to each other. Protein-protein interactions are complex and accurately predicting the conformations of protein-protein complex is challenging. Deep generative models may not fully capture the complexity of the protein-protein complex landscape, leading to incorrect conformational sampling. Deeply generative models rely heavily on training data, and the method of sampling conformations can affect the diversity and quality of the generated conformations. If the training dataset is biased towards some protein-protein complex conformations, or rare conformations can lead to incorrect generated conformational models. In addition, the MD simulations of protein-protein complex can be complemented with their conformational generation models to enhance the understanding of the system conformational dynamics of protein-protein complex.

**Figure 5.**
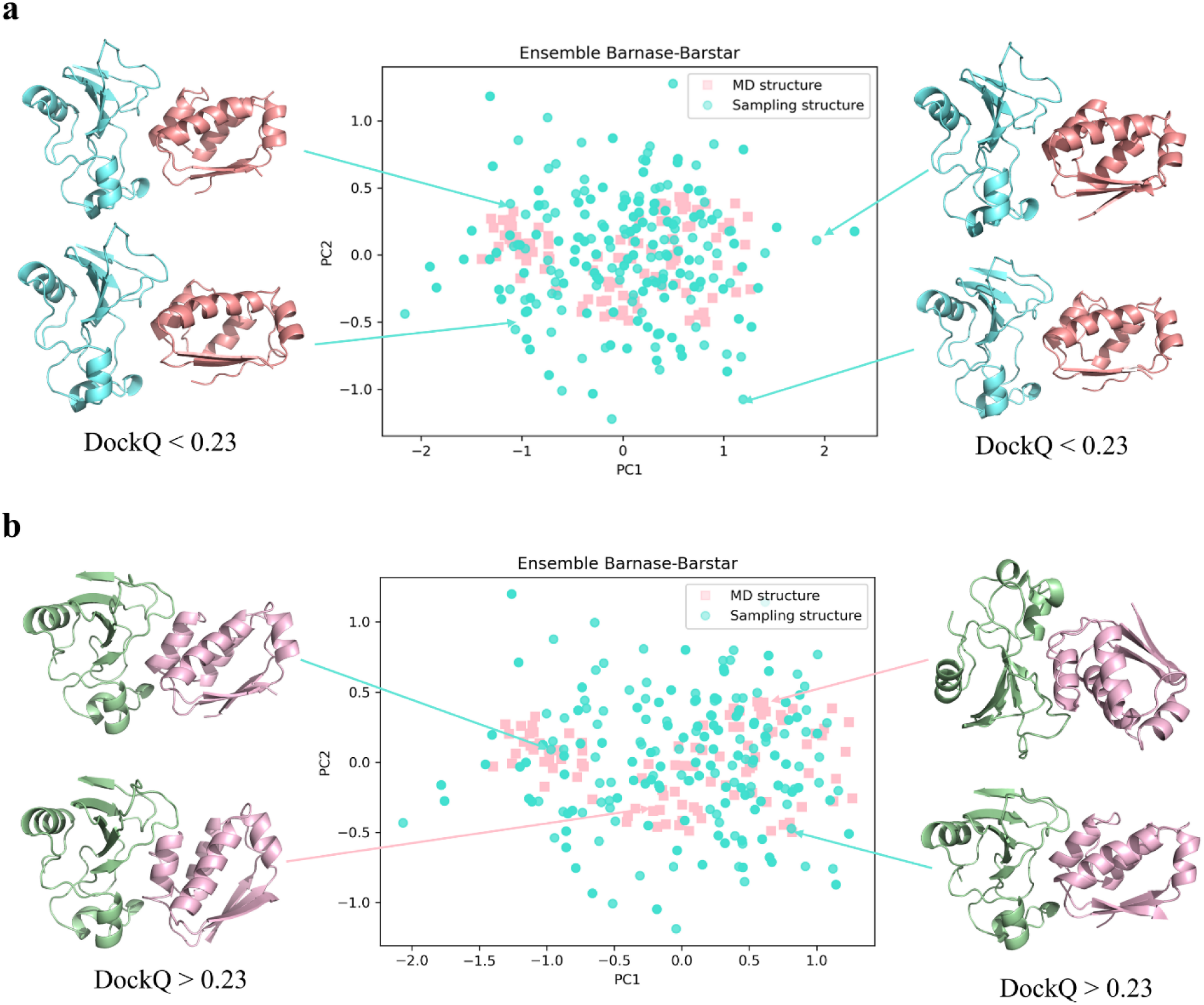
The PCA visualization of the conformational distributions of protein-protein complex ensembles: MD ensembles (color: pink) and sampling ensembles (color: greencyan). **a**, The PCA visualization of MD ensembles and all sampling ensembles and shows the generated conformational models with DockQ score < 0.23. **b**, The PCA visualization of MD ensembles and sampling ensembles with DockQ score ≥ 0.23 and shows representative conformational models. The distance of PCA is defined as a reduction from the Euclidean distance of the backbone residues of the crystal structure.

### Explainability Analysis

The knowledge of amino acid residues and residue pairs from different partners that are critical to the conformational, mechanical and dynamical mechanisms of protein-protein complexes is the basis for understanding many protein functions. The attention mechanism of AlphaPPImd captures the attention weights between key residues, and provides mechanistic insights into protein-protein binding^54–56^. The attention mechanism assigns weights to different residue pairs based on their importance in the conformational changes of protein-protein complex. Residue pairs with higher attention weights indicate stronger or more frequent interactions/effects, which are essential for stabilizing protein-protein complex. Protein-protein interactions are concentrated at the protein-protein interface, so residue pairs located at the protein-protein interface tend to have higher attention weights. Residue pairs with high attention weights at protein-protein interfaces are critical for specific recognition and binding between proteins, which is essential for understanding the stability and specificity of protein-protein complex. We explored the key residues and residue pairs between the two chains that have an impact on the dynamics and conformation of the barnase-barstar complex by attention weights. As shown in **Figure 6a**, different colors are applied with attention weights ranging from 0 to 1. The higher the intensity of red color has the higher attention weight for the particular residue pairs, indicating that its inherent amino acids contribute the most to the dynamics and conformation of the barnase-barstar complex, whereas the higher the intensity of blue color has the lower attention weight for the particular residue pairs whose inherent amino acids contribute the least to the dynamics and conformation of the barnase-barstar complex. We showed the residues with higher attention weights on the crystal structure of the barnase-barstar complex, as shown in **Figure 6b** (drawn by Pymol^57^). The results show that the key residues captured by the AlphaPPImd model are mainly located at the interface of protein-protein interactions, loop and helix. Residue pairs with high attention weights generally correspond to regions undergoing significant conformational changes or involved in key dynamic processes. It means that deep generative model captures key residues from the MD trajectories of the barnase-barstar complex that affect its dynamics and conformation, which can be used to complement the MD results.

**Figure 6.**
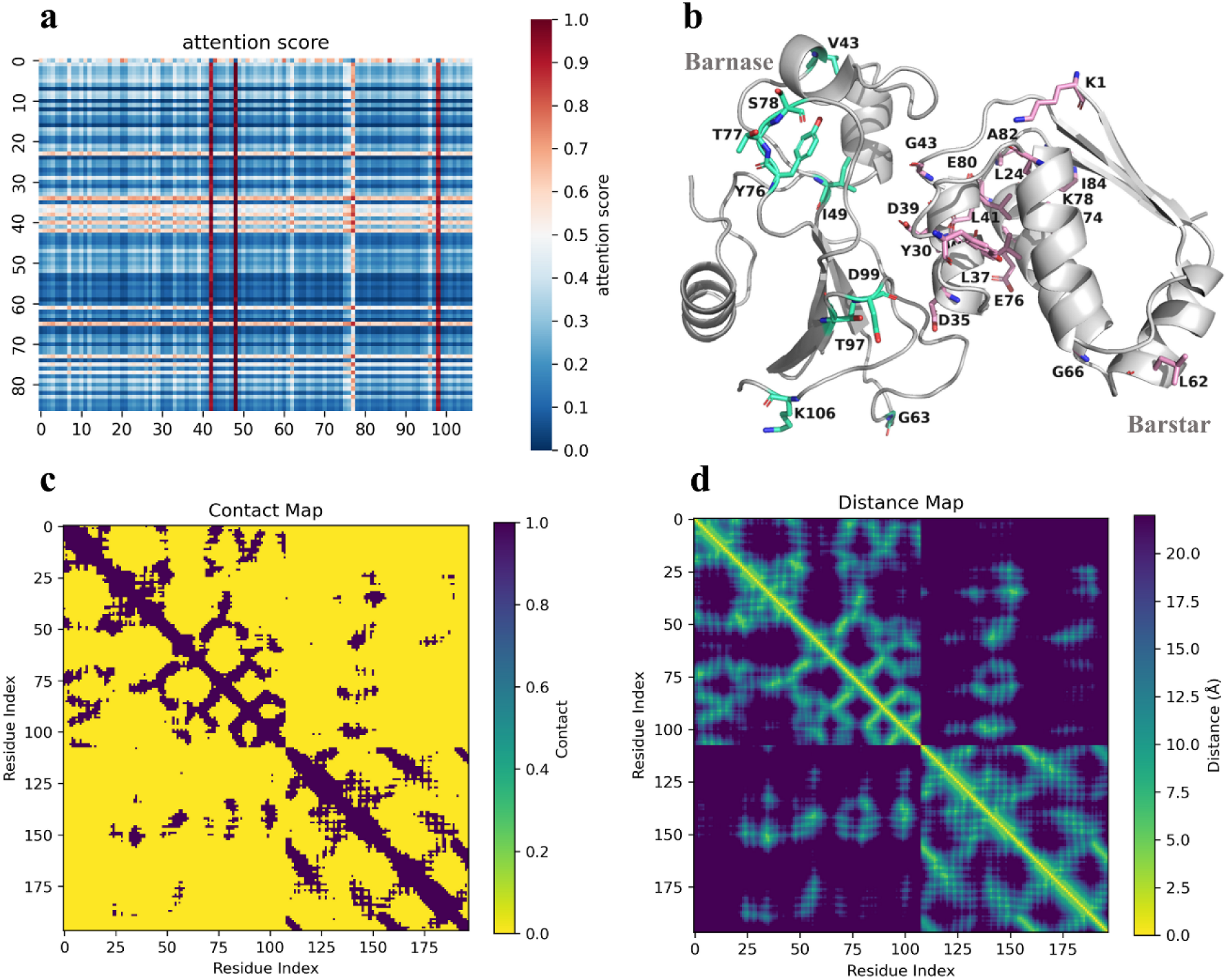
The explainability analysis. **a**, The attention score map of amino acid residue pairs between the two chains of the barnase-barstar complex. **b**, The mapping of attentional residues in the 3D structure of barnase-barstar complex. The key residues of barnase and barstar are highlighted in greencyan and pink stick, respectively. **c**, The contact map of the crystal structure of the barnase-barstar complex. The rows and columns of the contact map correspond to the residue index of the protein-protein complex. **d**, The distance map of the crystal structure of the barnase-barstar complex. The rows and columns of the distance map correspond to the residue index of the protein-protein complex.

The contact map is a binary two dimensional matrix representation of residue interactions in the protein structure. The rows and columns of the contact map correspond to the residue index of the protein-protein complex. The cells in contact map are usually colored to indicate the strength or frequency of the interaction. Darker colors may indicate stronger interactions or higher frequency of contacts, while lighter colors may indicate weaker interactions or fewer contacts. The diagonal lines of contact map represent intramolecular interactions within each protein. These interactions occur between residues within the same protein and are not involved in protein-protein interactions. The non-diagonal elements represent intermolecular interactions between residues of two proteins. These interactions are essential for stabilizing protein-protein complexes. The distance map is a two dimensional matrix representation of the distances between all possible residue pairs in the protein structure, where residue-residue distances are shown as a color gradations. The distance map illustrates the pairwise distances between residues in a protein-protein complex. The binding interface between two proteins is usually characterized by clusters of short distances between interacting residues. Shorter distances between residues imply stronger interactions. We calculated the contact map (cutoffs =13 Å) and distance map of the barnase-barstar complex, as shown in **Figure 6c and 6d**. We were able to observe that most of the key residue pairs between the two chains revealed by the contact/distance map, which appeared in residue pairs with higher attention weights. The continuous contact/distance maps of the MD trajectories of protein-protein complexes are taken as representations to train the deep generative model to generate novel conformational models, potentially producing higher resolution models of protein-protein complex.

### Independent conformational ensembles exploration of PPI complex

We selected MDM2-p53 protein-protein interaction to perform independent validation. The crystal structure of the MDM2-p53 complex (PDB ID: 1YCR) was used for the simulation system^58^, and the detailed MD protocol can be found in Supporting Information note B. The simulation time step was set to 2 fs, and then ten independent 30 ns MD production were performed. All simulations were performed by OpenMM 7.7^23^. The MD trajectories of the MDM2-p53 complex were randomly selected 1000 frames input to the model to generate 1000 novel basin-encoded frames, which were then reconstructed into novel conformational models of the MDM2-p53 complex.

We randomly selected a representative conformation with RMSD close to 2 Å from the 1000 generated conformational models of the MDM2-p53 complex. Then, the RMSD and DockQ scores were calculated by overlapping it with the crystal structure, as shown in **Figure 7a**. The 1000 frame conformational models generated by the AlphaPPImd model were filtered with DockQ < 0.23 (“incorrect”), and then PCA analysis was performed with conformational models extracted from the MD trajectories of the MDM2-p53 complex (**Figure 7b**). As shown in **Figure 7c**, different colors represent different attention weights. Residue pairs with stronger red color have higher attention weights, indicating that they contribute the most to the dynamics and conformation of the MDM2-p53 complex, while residue pairs with stronger green color have lower attention weights, and they contribute the least to the dynamics and conformation of the MDM2-p53 complex. As shown in **Figure 7d**, we show residues with higher attention weights on the crystal structure of the MDM2-p53 complex. The results showed that the key residues captured by the AlphaPPImd model are mainly located at the interface of MDM2-p53 interaction. Independent validation demonstrates the generalization of the model to other protein-protein complexes.

**Figure 7.**
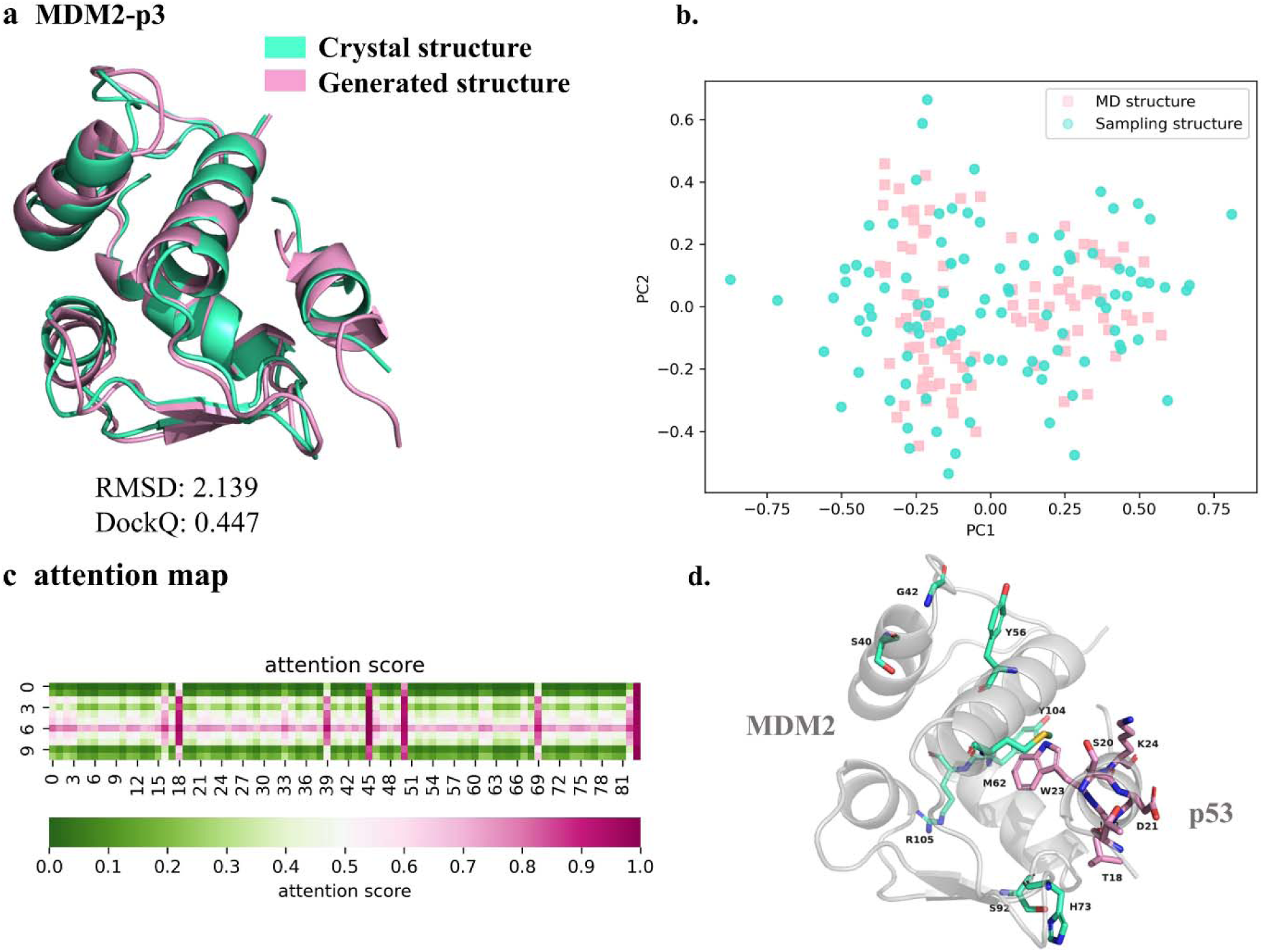
**a**, overlap of the generated conformational models of the MDM2-p53 complex (color: pink) with the crystal structure of the MDM2-p53 complex (color: greencyan), show the RMSD values and DockQ score of the generated conformational models with the crystal structure. **b**, The PCA visualization of MD ensembles and sampling ensembles with DockQ score ≥ 0.23 and shows representative conformational models. **c**, The attention score map of amino acid residue pairs between the two chains of the MDM2-p53 complex. **d**, The mapping of attentional residues in the 3D structure of MDM2-p53 complex. The key residues of MDM2 and p53 are highlighted in greencyan and pink stick, respectively.

## CONCLUSION

MD simulations are widely utilized to investigate the conformational ensembles and dynamics of protein-protein complexes, but there are significant limitations in sampling efficiency and computational cost. Deep generative models have achieved unprecedented advancements in the field of computational biology. Generative deep learning has been broadly used for protein conformational sampling. There is still a scarcity of applications of deep generative models to explore the conformational ensembles of protein-protein complexes.

In this study, we present a strategy for accelerated conformational sampling of protein-protein complex by combining Transformer-based generative model with MD simulation trajectories, which can extend protein-protein physical states that are difficult to discover by traditional MD simulation. AlphaPPImd can generate conformations beyond the MD time scale. The results of scoring the generated conformational models of the protein-protein complex showed that most of the generated conformational models were acceptable. The conformational ensembles analysis showed that the learned latent space could be used to generate unsampled conformations of protein-protein complex. The higher weights of residues and residue pairs captured by the attention mechanism were visualized. Deep generative mold learns critical residues from multiple MD trajectories that influence the conformational and dynamical mechanisms of protein-protein complexes, and provides mechanistic insights in protein-protein binding.

Although there are a few incorrect conformational models in the generated conformations, it generates more diverse conformations. The deep generative model generated many acceptable conformational models of protein-protein complex, but their quality was low compared to the physical conformation of MD. Training the deep generative model with continuous contact/distance maps of protein-protein complex MD trajectories as inputs or conditions may generate higher resolution conformational models of protein-protein complexes.

In the future, we can combine the rich conformational information of protein-protein complexes, the prediction of protein-protein interactions/binding affinity^59^, PPI binding site prediction^60^, the prediction of PPI-modulator interaction^61^, and the generative design of PPI modulators for accelerating the study of PPI targets and the design/screening of modulators^62,63^.

## ASSOCIATEDCONTENT

### Data Availability

The datasets and source code are publicly available at https://github.com/AspirinCode/AlphaPPImd.

### Supporting Information

Supporting Materials, analysis of molecular dynamics simulation of barnase-barstar complex, molecular dynamics simulation of MDM2–p53; Supporting Tables, Architecture of the Transformer neural networks; Supporting Figures, RMSD plot of molecular dynamics simulation of barnase-barstar complex.

## AUTHOR INFORMATION

### Authors

**Jianmin Wang** – The Interdisciplinary Graduate Program in Integrative Biotechnology, Yonsei University, Incheon 21983, Republic of Korea;

**Xun Wang** – School of Computer Science and Technology, China University of Petroleum, Qingdao, 266580, Shandong, China; High Performance Computer Research Center, University of Chinese Academy of Sciences, Beijing, 100190, China;

**Yanyi Chu**– Department of Pathology, Stanford University School of Medicine, Stanford, CA 94305, USA;

**Chunyan Li** – School of Informatics, Yunnan Normal University, Kunming, China;

**Xue Li** – School of Computer Science and Technology, China University of Petroleum, Qingdao, 266580, Shandong, China;

**Xiangyu Meng** – School of Computer Science and Technology, China University of Petroleum, Qingdao, 266580, Shandong, China;

**Yitian Fang**– School of Life Sciences and Biotechnology, Shanghai Jiao Tong University, Shanghai, 200240, China;

## AUTHOR CONTRIBUTIONS

J. Wang collected data, developed the model, analyzed the data, and wrote the manuscript; J. Mao and C. Li developed the model and analyzed the data; X. Wang, Y. Chu, X. Li, X. Meng and Y. Fan helped to refine the research through constructive discussions and revised the manuscript; K. No, J. Mao, X. Zeng supported and supervised the project, interpreted the results, and wrote revisions to the manuscript.

### Notes

Any additional relevant notes should be placed here.

## ACKNOWLEDGMENTS

This research was supported by the Yonsei University graduate school “Integrative Biotechnology”.

## Notes

### Competing Interest Statement

The authors have declared no competing interest.

### Summary of Updates

Figure 5 has also been revised. Figure 7 is corrected. We have revised novel in the section of evaluation metrics.

